# A pan-cancer catalogue of driver protein interaction interfaces

**DOI:** 10.1101/015883

**Authors:** Eduard Porta-Pardo, Thomas Hrabe, Adam Godzik

## Abstract

Despite their critical importance in maintaining the integrity of all cellular pathways, the specific role of mutations on protein-protein interaction (PPI) interfaces as cancer drivers, though known for some specific examples, has not been systematically studied. We analyzed missense somatic mutations in a pan-cancer cohort of 5,989 tumors from 23 projects of The Cancer Genome Atlas (TCGA) for enrichment on PPI interfaces using e-Driver, an algorithm to analyze the mutation pattern of specific protein regions such as PPI interfaces. We identified 128 PPI interfaces enriched in somatic cancer mutations. Our results support the notion that many mutations in well-established cancer driver genes, particularly those in critical network positions, act by altering PPI interfaces. Finally, focusing on individual interfaces we are also able to show how tumors driven by the same gene can have different behaviors, including patient outcomes, depending on whether specific interfaces are mutated or not.

## Introduction

Cancer patients are extremely heterogeneous in their response to treatments and outcomes. The first step towards the understanding of this variability was the identification of the multitude of genes that cause cancer, the so-called cancer driver genes^1^. In that sense, the completion of The Cancer Genome Atlas (TCGA) and other large-scale cancer genomics projects was a watershed event, as it provided the critical mass of data needed to identify driver alterations in most types of cancers^2–15^. Moreover, cancer types that previously were thought unique were found to represent different subtypes with different outcomes depending on the specific genes altered in each patient^16^. Since the start of the TCGA project, the catalogue of cancer drivers has increased and become more accurate^17^ thanks not only to the data generated by the project itself, but also to the development of multiple, complimentary algorithms that search for cancer driver genes using different approaches. Some of these methods identify cancer drivers by searching for genes with higher than expected mutation rates^18,19^, whereas others identify genes that tend to accumulate damaging mutations^20^ or protein regions with an unusually high proportion of mutations^21,22^.

Nevertheless, the catalogue of cancer driver genes is far from complete and because of extreme mutation diversity, it is hard to extend only by increasing the size of the datasets^19^. An complimentary approach towards that goal is to use other approaches that integrate cancer mutation profiles with other types of biological data to improve their analysis. For example, merging the mutation profile of cancer patients with biological networks can be used to find pathways and protein complexes that are recurrently mutated in cancer and are, therefore, likely drivers^23^. Note that these pathways and complexes can only be identified when adding the mutation profiles of all the components, because each individual protein is rarely mutated and missed by the standard approaches. Similarly, we can also include information on protein structure in our analysis to try to identify protein regions enriched in cancer mutations^22,24–26^. The underlying idea for this approach is that genes (and the proteins they encode) are not monolithic entities, but instead have different regions usually responsible for different functions. Such regions may include, for example, an enzymatic domain, a PPI domain or interface, or a phosphorylation site. In that context, it is possible that a given protein acts as a driver only when a specific region is mutated. Such fine grain approaches are not only capable of finding novel cancer drivers, but they also can help explain some of the variability between tumors or cancer cell lines apparently driven by the same gene^27^. We have previously developed an algorithm, e-Driver, which exploits this feature to identify cancer driver genes based on linear annotations of biological regions such as protein domains^22^. Despite encouraging results, the algorithm still had some limitations, as many features, such as epitopes or interaction interfaces, may be discontinuous at the sequence level and could not be analyzed.

Here we introduce an extended version of e-Driver that uses information on three-dimensional structures of the mutated proteins to identify specific structural features. Then, the algorithm analyzes whether these features are enriched in cancer somatic mutations and, therefore, are candidate drivers. While technically the analysis can be applied to any structural feature or region, we focused our attention to protein-protein interaction (PPI) interfaces. Many known cancer driver genes are located in critical regions of the PPI network (interactome), usually in network hubs or bottlenecks^28^. Moreover, many cancer somatic alterations, including passenger mutations, alter PPI interfaces, either destroying existing interactions or creating new ones^29,30^. Last but not least, despite their critical importance in every cellular function, the specific role of PPI interfaces as potential cancer drivers has never been systematically analyzed.

Our analysis identified PPI interfaces in 128 genes (interface driver genes), including 28 well-known cancer driver genes, which are strongly enriched in somatic missense mutations. The role of the remaining 100 genes as cancer drivers will, obviously, have to be verified experimentally, though we find some attributes that suggest that some of them are indeed true drivers. For example, all the interface driver genes (including the 28 known drivers) have an unusually high number of interactions, not only when compared to the rest of the genes in the interactome, but also when compared to other cancer driver genes. We also identified numerous interfaces in genes related to the immune response, particularly in HLA-like and complement molecules. The role of the immune system in cancer treatment and evolution is gaining attention and we provide new details regarding which interactions seem to be most affected by somatic mutations. Finally, we show how, depending on which interface or protein region is altered, tumors thought to be driven by the same cancer gene might have radically different behaviors and patient outcomes.

## Results

### e-Driver reveals driver interfaces

We assembled a data set consisting of 5,989 tumors from 23 cancer types from The Cancer Genome Atlas^31^ (Supplementary Table 1). The number of samples per tumor type ranged from 56 for uterine carcinosarcoma, to 975 for breast adenocarcinoma (Supplementary Figure 1). Consistent with previous reports^32^, the average number of missense mutations per sample is highly variable among cancer types (Supplementary Figure 2), with melanoma having the highest (429 missense mutations per sample) and thyroid carcinoma the lowest (11 missense mutations per sample).

**Figure 1.**
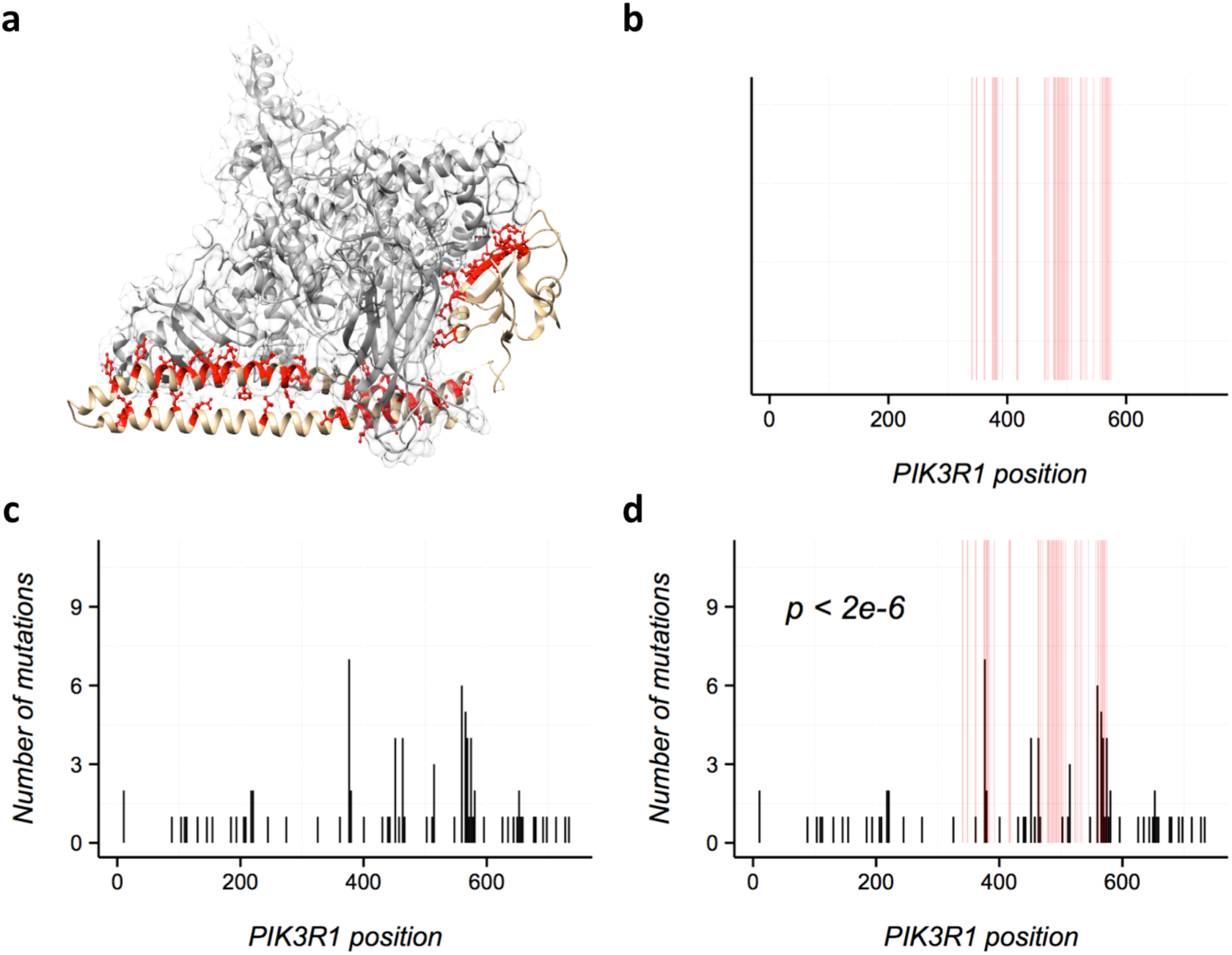
Using e-Driver to analyze PIK3R1 PPI interfaces. a) The PDB coordinate set 3HMM contains 2 chains, A (a region of PIK3CA, shown in gray) and B (a region of PIK3R1, shown in brown). Residues from PIK3R1 colored in red make the interaction interface with PIK3CA. b) Using BLAST we then map these residues to the corresponding ENSEMBL protein, define a PPI interface (shown in red). Note that the interface is not continuous in sequence. c) Distribution of mutations in PIK3R1 across all cancer types. d) Using e-Driver we can identify the interface between PIK3R1 and PIK3CA as strongly enriched in missense somatic mutations (p < 2e-6).

**Figure 2.**
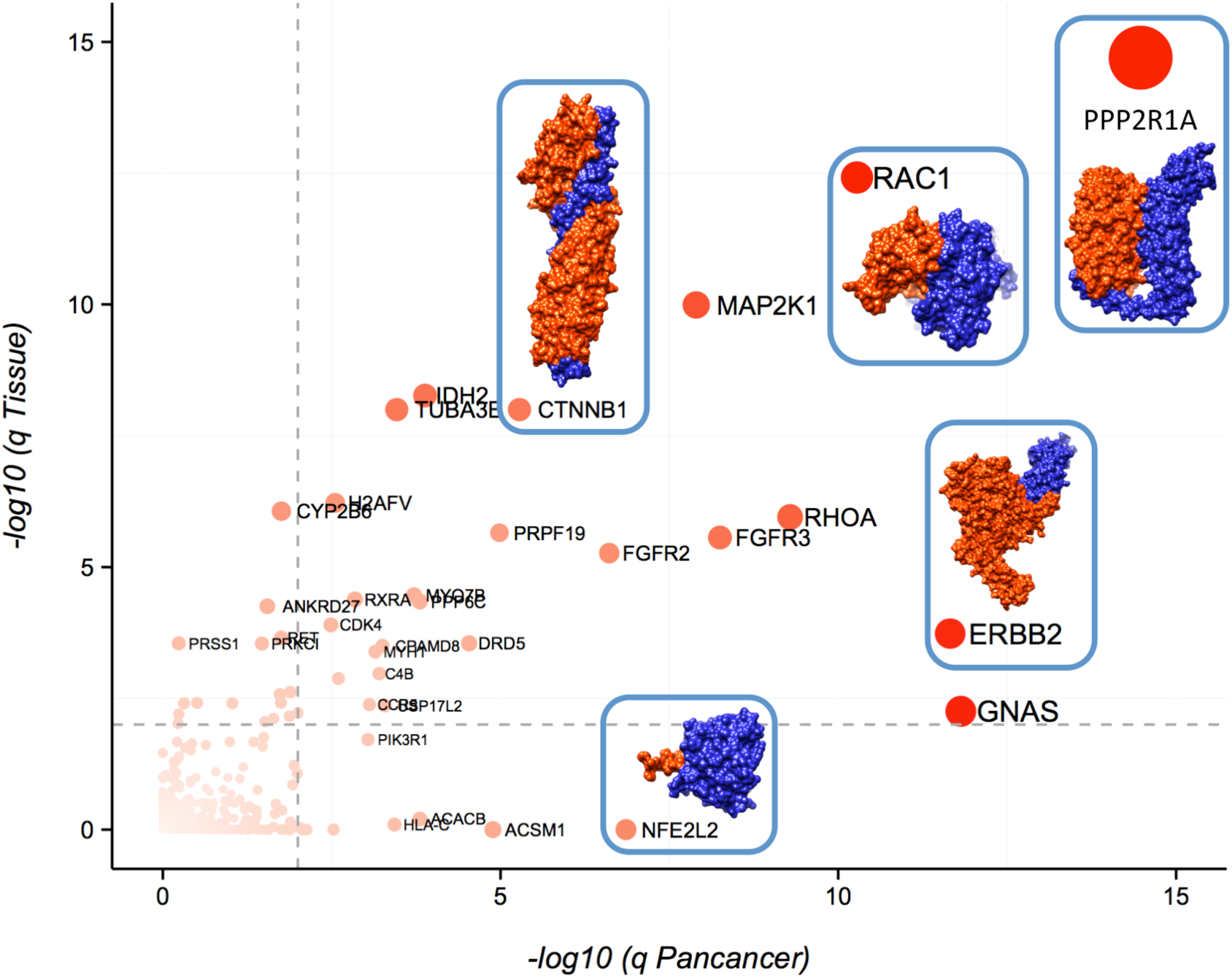
Genes with driver interfaces. X,Y coordinates reflect the q value (FDR) in the Pancancer analysis and their lowest q value in all 23 project-specific analysis, respectively. Gray dashed lines are located at 0.01 FDR for reference. Dots are colored, sized and labeled according to their FDR: genes with an FDR of 1 are colored in white, are smaller and have no label whereas genes with lower FDRs are more red, bigger and labeled. Note that there are 8 genes that are not in the plot because their FDR value was too small (FDR < 1e-15): *TP53, EGFR, KRAS, HRAS, NRAS, RPSAP58, PIK3CA* and *UBBP4.* Some driver interfaces are also illustrated in the plot, with the chain belonging to the gene with the driver interface colored in orange and the interacting chain colored in blue. The PDB coordinates used are 3IFQ for CTNNB1, 1S78 for ERBB2, 2FLU for NFE2L2, 3B13 for RAC1 and 3FGA for PP2R1A.

We then compiled a list of currently known, high-confidence PPI interfaces using 18,651 protein structures downloaded from PDB (Online methods). In short, we defined a PPI interface as the set of residues from a given chain that are within 5 angstroms of any residue from a different chain in the same set of PDB coordinates (Figure 1a). We identified 122,326 different PPI interfaces between 70,199 PDB chains (Online methods). Finally, we used BLAST to map the residues from the PDB datasets to gene sequences in the ENSEMBL human genome. Overall we mapped the PDB coordinates to 11,154 protein isoforms in 10,028 different human genes. The mapping covers roughly 30% of the length of human proteome, with 6% of the proteome being mapped to at least one PPI interface.

Mutations from all cancer datasets (n = 868,508) seem to be distributed randomly across the proteome, with approximately 30% of mutations (n = 285,942) being in regions mapped to structures and around 6% in PPI interfaces (n = 67,174). However, in the case of known cancer driver genes ^1,17^, regions covered by structures have between two and three times more missense mutations than expected by chance (Supplementary Figures 3-4). This enrichment, while variable and dependent on the cancer type, is particularly high in regions involved in the PPI interfaces. For example, PPI interfaces from cancer driver genes have more than tree times as many mutations as would be expected by chance in breast adenocarcinoma, glioblastoma lower grade glioma rectal adenocarcinoma or uterine carcinosarcoma (Supplementary Figure 4) confirming that, indeed, PPI interfaces might play key roles in carcinogenesis.

**Figure 3.**
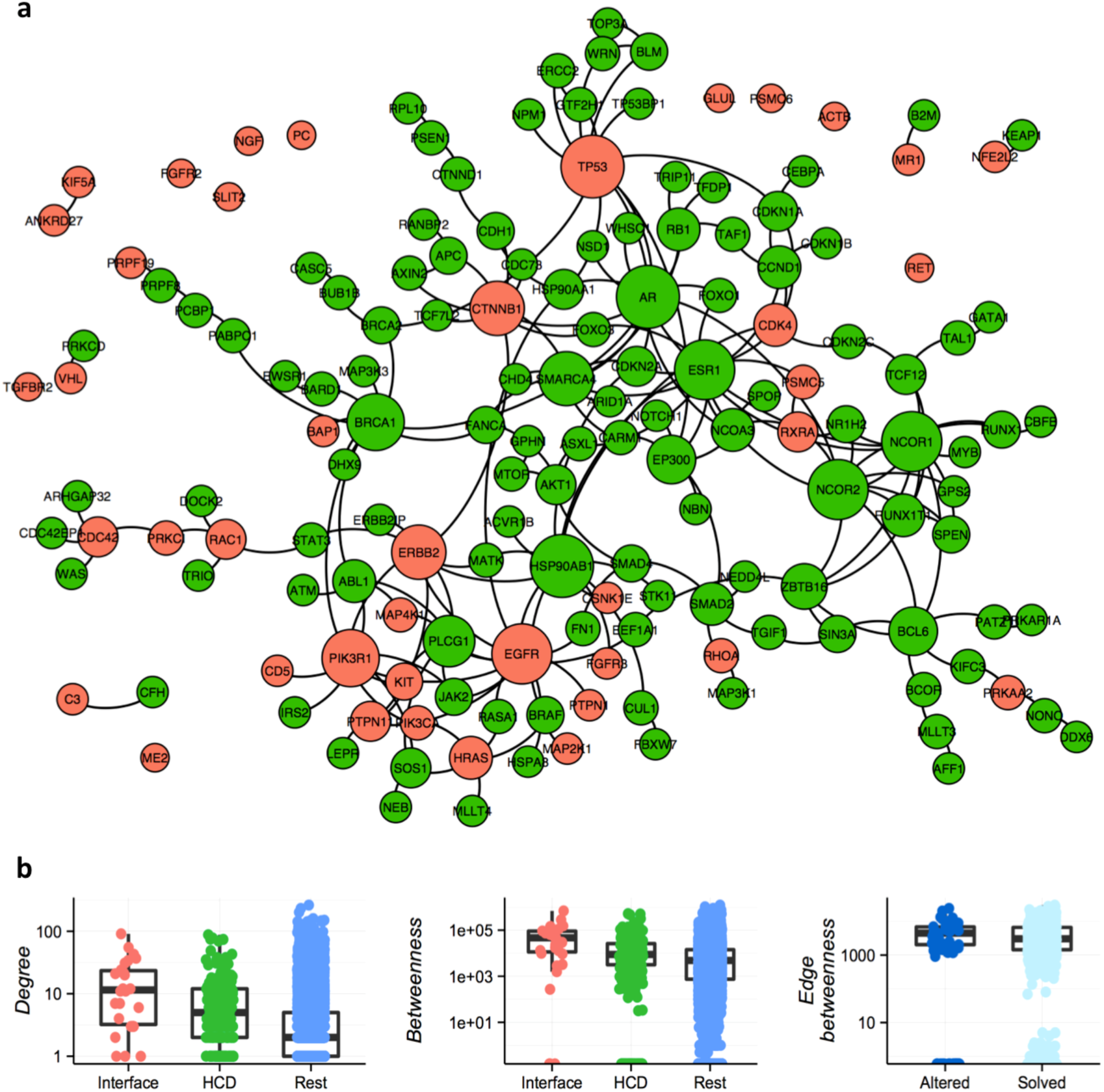
Network properties of interface-driver genes. a) Most genes identified by e-Driver3D (shown in red) cluster together with other cancer driver genes (shown in green). Genes are sized according to their number of interactions (genes with more interactions are bigger and *vice versa*). b) Interface driver genes are critical elements of the interactome. They have more interactions (higher degree, left panel) and betweenness (center panel) than other cancer driver genes or the rest of the genes in the interactome. Moreover, interactions between interface-driver genes (shown in dark blue, right panel) are more central (higher edge betweenness) than other structurally solved interactions (shown in light blue).

**Figure 4.**
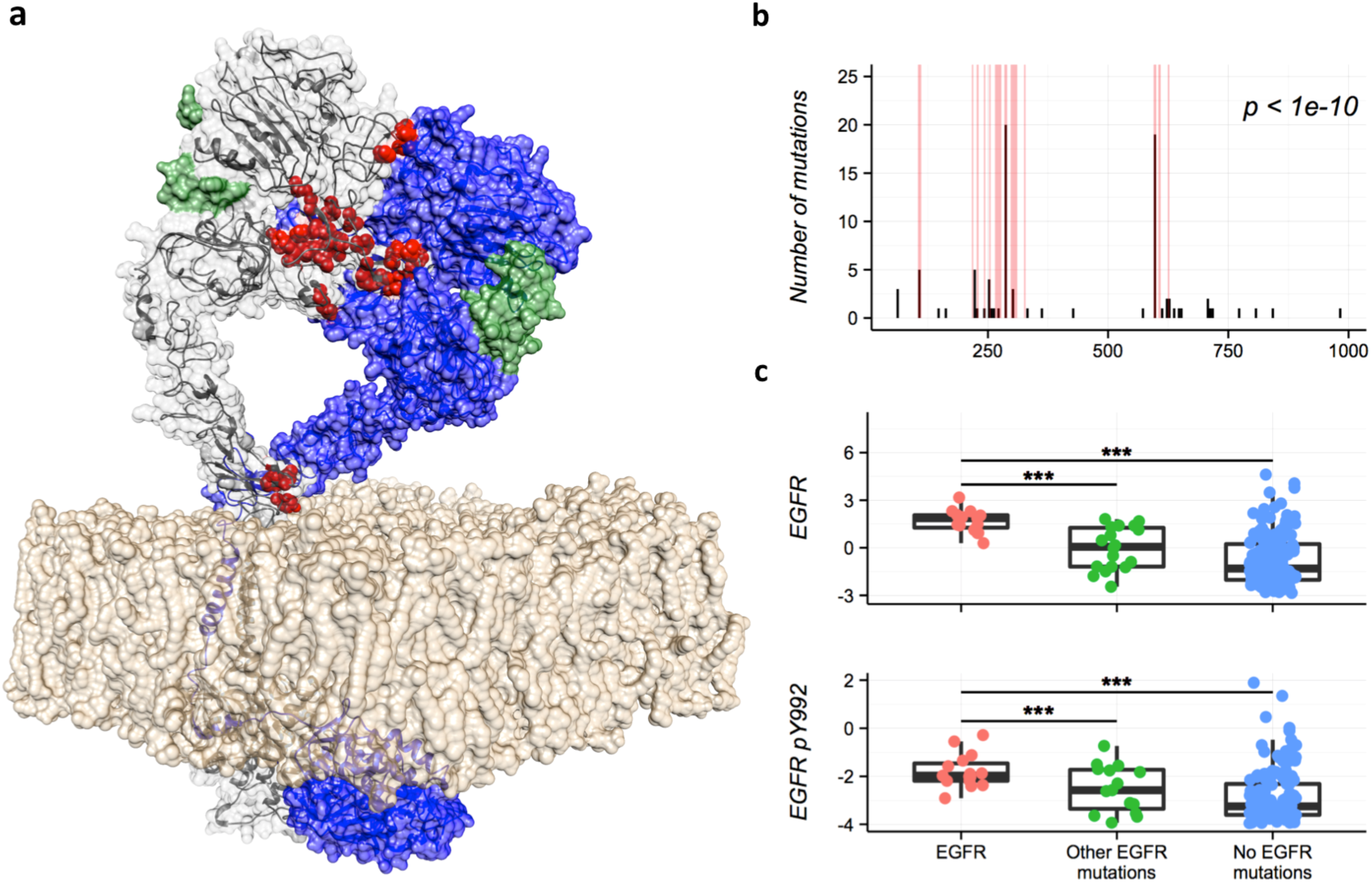
Protein expression changes induced by mutations in EGFR dimerization interface. a) Structure showing an EGFR dimer in active conformation (based on^41^). The residues involved in the EGFR-EGFR interaction for the left EGFR molecule are shown in red, the other EGFR protein is shown in blue, the two EGF ligands are shown in green and the lipid bilayer in brown. b) Histogram showing that most glioblastoma mutations (black bars) in EGFR are located in its dimerization interface (red bars). c) Protein expression in different glioblastoma populations. Patients with mutations in EGFR’s dimerization interface (shown in red) have higher levels of EGFR and phosphorylated EGFR proteins than those with other or no EGFR mutations (shon in green and blue respectively).

Next, we used e-Driver to analyze individual PPI interfaces in each of the 23 individual cancer projects, as well as in the Pan-cancer dataset consisting of the combination of all of them. Briefly, e-Driver compares the observed number of mutations in a protein region with the expected value according to the length of the region and the length of the protein. We had previously used e-Driver to analyze the distribution of cancer somatic mutations in PFAM domains and intrinsically disordered regions and showed, for example, that different domains in the same protein can be driving different types of cancer. Here, we adapted e-Driver to analyze discontinuous features, such as PPI interfaces derived from 3D structures. The whole process is exemplified in Figure 1 for PIK3R1 and its interaction interface with PIK3CA.

We identified a total of 128 interface driver genes in either one of the cancer projects or in the Pan-cancer analysis (Figure 3, Supplementary Tables 3-26). There is significant overlap between the genes identified in this analysis and lists of known cancer genes. For example, 28 interface driver genes (22%) are included in a list of high-confidence driver genes derived from previous analyses of TCGA data^17^ (p < 1e-10, odds ratio 10), and 26 are part of the Cancer Gene Census^1^ (p < 1e-9, odds ratio 5). Note that our predicted interface driver genes do not include 268 and 476 genes belonging to the TCGA high confidence driver genes list and CGC respectively. These might not have been identified in our analysis not only because they do not have driver interfaces, but also because we currently do not have a structure with a PPI interface to match to them and, thus, they were not included our analysis. For example, we only have a structure for 50% of all the mutations in known driver genes, the rest might also be altering interactions, but we cannot know it until we increase the structural coverage of the human proteome.

Some of the driver interfaces identified here contain known cancer hotspots. For example, *NFE2L2,* a gene involved in cancer progression and drug resistance, is usually activated by mutations that disrupt the interaction with its repressor *KEAP1.* We mapped 36 mutations from *NFE2L2* to the structure showing its interaction with its repressor *KEAP1* (PDB 2FLU, shown in Figure 2). In agreement with previous observations^15^, all but two of the mutations (94%) in *NFE2L2* involve interface residues, likely disrupting the interaction between the two proteins and activating *NFE2L2.*

Our results also highlight similarities and differences across related driver genes. For example, receptor tyrosine kinases, particularly members of the ERBB and FGFR families, are mutated in many cancer samples and frequently act as drivers. We found two ERBB proteins among the interface driver genes, ERBB2 and EGFR. These two proteins are both strongly enriched in mutations in their dimerization interfaces, while the ligand-binding region is rarely mutated (Supplementary Figure 5). We also identified two proteins from the FGFR family: FGFR2 and FGFR3. Again, these two proteins have similar mutation profiles, with both proteins having most of their missense mutations in the region that interacts with the ligand (Supplementary Figure 5), while leaving the dimerization interface intact. This, however, contrasts with the mutation pattern of the ERBB receptors, where, as we have explained, the ligand-binding region is rarely mutated. Since some of the most successful therapeutic antibodies against EGFR target the dimerization interface identified by our method, it is possible that antibodies against FGF receptors need to target the ligand-binding region in order to be successful^33^.

**Figure 5.**
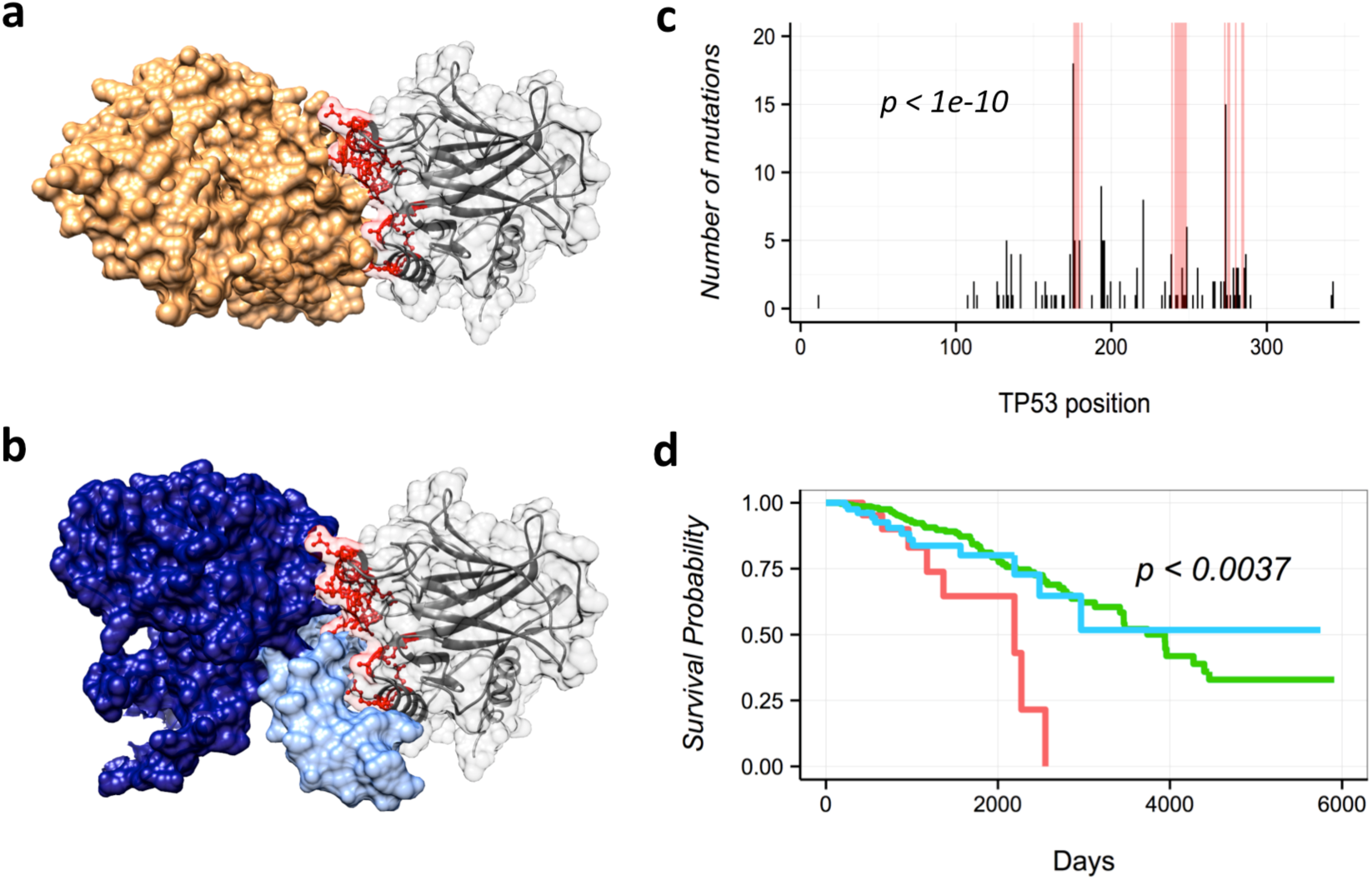
Mutations in TP53 interface predict patient’s outcomes. a) Our analysis identified the interface between TP53 and SV40 (shown in gray and orange respectively) as strongly enriched in mutations (panel c). b) This interface is the same as the one that TP53 uses to interact with other TP53 molecules (shown in dark blue) and DNA (shown in light blue). d) Breast cancer patients with mutations in this interface (shown in red) have worse outcomes than those with other or no TP53 mutations (shown in blue and red respectively).

Among the 100 interface driver genes that are not currently classified as cancer drivers (Supplementary Table 2), we find several cases where literature and biological evidence support their role in cancer. This is the case, for example, for CDC42. This protein is a small GTPase, involved in cellular functions key to cancer progression, such as cell migration and mitosis^34^. Moreover, it also interacts with 4 known cancer driver genes (ARGHAP32, WAS, PRKC1 and CDC42BP1), providing more evidence as to its role as a cancer driver. Another subset of these 100 potential new cancer driver genes have functions related to immunity. Given the growing body of evidence showing that the immune system plays a key role in cancer progression and patients outcomes^35,36^, we analyzed these interfaces in more detail to try to find novel insights about the interplay between tumors and immune cells. For example, a recent pan-cancer analysis identified a subnetwork of proteins around HLA class I as being recurrently mutated in cancer^23^. Our analysis also identified several antigen-presenting molecules as potential cancer drivers, including one class I (HLA-C), one class II (HLA-DRB1), and three HLA-like proteins (CD1C, CD1E and MR1). Note, also, that HLA-C has been recently identified as a likely driver in head and neck cancer^15^. Another interesting group of immune-related proteins that identified in our analysis include several elements of the complement cascade (C3, C4B and C5) or complement regulators and inhibitors (CFHR4, CFI and CPAMD8). The complement molecules C3 and C4 have been previously associated with cancer progression and activation of PI3K signaling^37^, whereas C5a is suspected to inhibit CD8 lymphocytes and natural killer (NK) cells, a subset of immune cells involved in the immune response towards tumors^38^.

### Interface driver genes are network hubs

Cancer driver genes are known to occupy critical positions in the interactome, as well as having more interactions and higher betweenness than the average gene^28^. Since our method identifies genes strongly enriched in mutations in their PPI interfaces, we hypothesized that they would also be located in these central positions of the protein interaction network. To test this hypothesis, we used a PPI network resulting from the merge of an unbiased experimental interactome and a curated network from the literature^39^. In that network, interface driver genes tend to interact between themselves more than expected by chance (p < 0.0001, Supplementary Figures 6-7), though this seems to be a general property of cancer driver genes. Moreover, interface driver genes have higher degree (number of overall interactions) and betweenness than the average gene (Figure 3), supporting our hypothesis. Remarkably, this is also true if we compare them to known cancer drivers with no driver interfaces. These results are consistent with the hypothesis that the main driver mechanism of the genes identified in our analysis is the alteration of their interaction interfaces.

### Consequences of mutations in driver interfaces

Even if the genes that we identified have more mutations than expected in some of their PPI interfaces, there are tumor samples with mutations in other regions of the same genes. With that in mind, we wondered if there are consistent differences between cancer samples belonging to each of these two groups. To explore this issue, we first used proteomics data^40^ and compared the expression levels of different proteins in tumors with mutations in the predicted driver interfaces with that of tumors with mutations in other regions of the same gene. To limit the impact of intrinsic tissue-variability in the protein expression levels, we limited our analysis to tissue-specific driver interfaces.

Though we could not analyze most of the interfaces due to lack of statistical power (there were not enough samples with proteomics data in both groups), we did find some interface-specific protein changes. For example, glioblastoma samples with mutations in EGFR’s dimerization interface have higher levels of both EGFR and phosphorylated EGFR (Y992 and Y1173) proteins than patients with other EGFR mutations (Figure 4), suggesting that EGFR signaling is stronger in these patients. Note that these results also agree with the hypothesis that the main molecular mechanism driving cancer in these genes is the disruption of certain interactions, as the cancer cells have different signaling levels depending on whether the gene is mutated in the identified driver interfaces or in another region.

Another example of interface-specific protein expression changes comes from TP53 and its interface with SV40 (Figure 5). Note that this interface is the same as the one that TP53 uses to dimerize and to bind to DNA (Figure 5b). Patients from eight different cancer types (bladder, breast, colon, endometrial, glioma, stomach, lung and head and neck) with mutations in this interface had significantly higher levels of TP53 protein than those with other or no TP53 mutations (Supplementary Figure 8). Moreover, patients with breast cancer had significantly worse outcomes (Figure 5d), suggesting that these mutations are more aggressive than other mutations in TP53 and that maybe different therapeutical approaches are needed in these cases. Note that traditional gene-centric analyses or the previous version of e-Driver cannot find these differences among patient subpopulations.

## Discussion

Here we explored the role of mutations in PPI interfaces as cancer driver events using our e-Driver algorithm and the mutation profiles of 5,989 tumor samples from 23 different cancer types. Though the interaction interfaces of some cancer driver genes have been studied before^42,43^, this is the first time that three-dimensional protein features, such as PPI interfaces, have been systematically used to identify driver genes across large cancer datasets. Previous large scale analyses, including our own, are either limited to linear features^22,24,44^, or do not define functional regions in three-dimensional structures but, instead, identify *de novo* three-dimensional clusters of mutations^25^. Our analysis identified several driver PPI interfaces in known cancer driver genes, such as TP53, HRAS, PIK3CA or EGFR, proving that our method can find relevant genes and that alteration of interaction interfaces is a common pathogenic mechanism of cancer somatic mutations. In fact, we found that cancer driver genes, as a group, are strongly enriched (over two-fold in most cancer types) in mutations in their PPI interfaces. Moreover, there is a strong correlation between the fact that a cancer driver gene is recurrently mutated in its PPI interfaces and how critical it is to the stability of the interactome in terms of both number of interactions and network betweenness. We also identified a series of driver interfaces in genes that are currently not known as cancer drivers. Some of these genes interact with known cancer drivers or are related to key cancer functions, such as the immune system, suggesting that they are, indeed relevant to carcinogenesis. Analysis of other genes with cancer driver interfaces is ongoing and we provide a complete list of such genes in the supplementary materials, as well as in our on-line resource Cancer3D^45^.

It is important to note that the analysis presented here was limited to high quality interfaces, predicted either from solved structures or from close homology models. However, about 70% of human proteome currently has no structural coverage. This fraction of the proteome includes both low complexity or disordered regions, and protein regions without available templates to model their 3D structures. Also, structures of many complexes are still unknown. In these cases, even if we know the structures of the subunits, we cannot define the PPI interfaces and these proteins are excluded from our analysis. Finally, even though we did not explore this issue here, there are other mutations that can have an impact in PPI interfaces, such as inframe indels or silent mutations^46^. Therefore, the results presented here represent only the tip of the iceberg of what can be achieved by including structural data in the analysis of cancer mutation profiles. We expect that our method will improve not only as more cancer genomes are added to existing repositories (increasing the statistical power of the analysis), but also as the structural coverage of the human proteome increases. We expect such increase to come from both new experimentally determined structures in public databases and the use of improved modeling tools^47,48^.

We also found that tumors apparently driven by the same gene can have surprisingly different behavior and outcomes depending on the specific PPI interface altered. This adds to a growing body of evidence suggesting that the current gene-centric paradigm in biology, while accurate in some cases, will not probably be enough to explain the complex genotype-phenotype relationships^49–53^. In the case of cancer, for example, it is known that the two most common mutations in PIK3CA, E545K and H1047L, contribute to carcinogenesis through different mechanisms^42^. The same is true for different types of mutations in KRAS ^54^ or, as we have shown here, for mutations in EGFR or TP53 among many others. All of the above suggests that, in order to predict the outcome of a patient or the best treatment option we will need to have more detailed knowledge about the consequences of a specific mutation than just the cancer driver gene where it is located. Such increase in detail and knowledge should include, not only information about the protein domain or PPI interface of the gene being altered, but also data about mutations in other regions of the network, as these can also influence the phenotype of a driver gene through synthetic interations^55^.

## Methods

All the raw data and algorithms used in this manuscript, as well as the results presented, can be downloaded from http://github.com/eduardporta/e-Driver. All the statistical calculations were done using R 3.1.0.

### Mutation data

We downloaded level 3 mutation data from the TCGA data portal (https://tcga-data.nci.nih.gov) for 5,989 tumor samples that belong to 23 different cancer types (Supplementary Table 1). We then used the Variant Effect Predictor tool to derive the consequences of each mutation in the different protein isoforms where it mapped^56^. We used gene and protein annotations from ENSEMBL version 72. We identified a total of 868,508 missense mutations in 19,196 proteins. Note that we only analyzed the longest isoform of each gene in order to minimize problems related to multiple testing.

### Identification of the protein-protein interfaces

We identified 18,651 protein structures with multiple chains in PDB (as of May 2014). Then, we analyzed all such structures to find the residues implicated in PPI interfaces. To that end, we defined a protein-protein interface in a chain as all the residues with a heavy atom within 5 angstroms of another heavy atom from a different chain, an intermediate value between the 4 and 6 angstroms seen in other references^30,57^. If a chain was in contact with multiple other chains, we defined a different interface for each chain-chain pair. Note that any specific interface does not have to be linear in sequence and that the same residue can be involved in multiple interfaces from different structures.

### Structure mapping

The mapping between ENSEMBL and PDB is the same as the one used in Cancer3D. Briefly, we queried the full PDB (March 2014), including non-human proteins, with every protein from ENSEMBL using BLAST. Every time we identified a PDB-ENSEMBL pair with an e-value below 1e^-6^ we used the BLAST output to map the residues from the ENSEMBL sequence to the PDB structure^45^.

### e-Driver analysis

We used e-Driver^22^ to identify interfaces that are enriched in somatic missense mutations. The algorithm calculates the statistical significance of deviation from the null hypothesis that the mutations are distributed randomly across the protein using a right-sided binomial test:

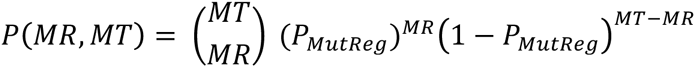

Where “P_MutReg_” is the ratio between the number of residues involved in the interface and the number of aminoacids in the entire protein, “M_R_” is the number of mutations in the interface and “M_T_” is the total number of mutations in the protein. Since it is possible that only a fraction of the protein is covered by the structure, we adjusted the algorithm to limit all the parameters to the structure-mapped region of the protein (for example “M_T_” refers to the total number of mutations in the region of the protein covered by the specific structure being analyzed, not the absolute total of mutations in the protein). The final step consists in correcting all the p values for multiple testing using the Benjamini-Hochberg algorithm. We considered as positives all of the interfaces with a q value below

### PPI Network analysis

Given the large number of available human PPI networks and the variability in their quality, we decided to use data from two recently published high-quality networks^39^. The first one consists in a set of 11,621 protein-protein interactions derived from the literature. In order to be in that network, an interaction must have been reported at least twice. The second network, described in the same publication, contains 13,945 interactions validated by, at least, two different experimental techniques. In our analysis, we used the network derived from the merge of these two, which contains 21,509 interactions between 7,463 proteins (Supplementary Table 27). We calculated the different network properties (node degree and node and edge betweenness) using the R package “iGraph”.

Finally, in order to find whether interface driver genes interact between themselves more than expected we compared the observed number of interactions in the actual network with that of 2,000 randomized networks. To estimate the distribution of z-scores, we repeated this procedure 100 times and calculated the z-score in each simulation. We also did this with known cancer driver genes from TCGA and CGC. A set of 300 genes was picked at random from the network each of the 100 times to be used as additional control.

### Protein expression and clinical data analysis

We downloaded level 3 clinical and protein expression data, whenever it was available, from the TCGA data portal. Then, for each statistically significant interface, we classified each sample into one of three groups: samples with mutations in the interface, samples with mutations in other regions of the same protein and samples with no mutations in that protein.

Finally, in the case of proteomics data, we used a two-sided Wilcoxon test to identify proteins with statistically significant differences between the first group and the other two. As for the clinical data, we used the Cox proportional hazards model, from the R package “survival”, to estimate whether mutation of a specific interface was a predictive feature for survival (p < 0.01) after correcting by age.

## Author’s contributions

EP-P and TH did the experiments. EP-P and AG concieved the experiments, interpreted the data and wrote the manuscript.

## Acknowledgements

The authors would like to thank the TCGA community for their great efforts and making all the data available. Funding for this research was provided by the Human Frontiers Science Program grant [grant number RGP0027/2011] and by the National Institutes of Health [grant number R01 GM101457].

## References

1. Futreal, P.A. et al. A census of human cancer genes. Nat Rev Cancer 4, 177–83 (2004).

2. Cancer Genome Atlas Research, N. Comprehensive genomic characterization of squamous cell lung cancers. Nature 489, 519–25 (2012).

3. Cancer Genome Atlas, N. Comprehensive molecular portraits of human breast tumours. Nature 490, 61–70 (2012).

4. Cancer Genome Atlas, N. Comprehensive molecular characterization of human colon and rectal cancer. Nature 487, 330–7 (2012).

5. Cancer Genome Atlas Research, N. Integrated genomic analyses of ovarian carcinoma. Nature 474, 609–15 (2011).

6. International Cancer Genome, C. et al. International network of cancer genome projects. Nature 464, 993–8 (2010).

7. Cancer Genome Atlas Research, N. Comprehensive genomic characterization defines human glioblastoma genes and core pathways. Nature 455, 1061–8 (2008).

8. Cancer Genome Atlas Research, N. Comprehensive molecular characterization of clear cell renal cell carcinoma. Nature 499, 43–9 (2013).

9. Cancer Genome Atlas Research, N. Genomic and epigenomic landscapes of adult de novo acute myeloid leukemia. N Engl J Med 368, 2059–74 (2013).

10. Davis, C.F. et al. The somatic genomic landscape of chromophobe renal cell carcinoma. Cancer Cell 26, 319–30 (2014).

11. Cancer Genome Atlas Research, N. Comprehensive molecular profiling of lung adenocarcinoma. Nature 511, 543–50 (2014).

12. Cancer Genome Atlas Research, N. Integrated genomic characterization of papillary thyroid carcinoma. Cell 159, 676–90 (2014).

13. Cancer Genome Atlas Research, N. Comprehensive molecular characterization of gastric adenocarcinoma. Nature 513, 202–9 (2014).

14. Cancer Genome Atlas Research, N. Comprehensive molecular characterization of urothelial bladder carcinoma. Nature 507, 315–22 (2014).

15. Cancer Genome Atlas, N. Comprehensive genomic characterization of head and neck squamous cell carcinomas. Nature 517, 576–82 (2015).

16. Hoadley, K.A. et al. Multiplatform analysis of 12 cancer types reveals molecular classification within and across tissues of origin. Cell 158, 929–44 (2014).

17. Tamborero, D. et al. Comprehensive identification of mutational cancer driver genes across 12 tumor types. Sci Rep 3, 2650 (2013).

18. Dees, N.D. et al. MuSiC: identifying mutational significance in cancer genomes. Genome Res 22, 1589–98 (2012).

19. Lawrence, M.S. et al. Discovery and saturation analysis of cancer genes across 21 tumour types. Nature 505, 495–501 (2014).

20. Gonzalez-Perez, A. & Lopez-Bigas, N. Functional impact bias reveals cancer drivers. Nucleic Acids Res 40, e169 (2012).

21. Tamborero, D., Gonzalez-Perez, A. & Lopez-Bigas, N. OncodriveCLUST: exploiting the positional clustering of somatic mutations to identify cancer genes. Bioinformatics 29, 2238–44 (2013).

22. Porta-Pardo, E. & Godzik, A. e-Driver: a novel method to identify protein regions driving cancer. Bioinformatics 30, 3109–14 (2014).

23. Leiserson, M.D. et al. Pan-cancer network analysis identifies combinations of rare somatic mutations across pathways and protein complexes. Nat Genet (2014).

24. Reimand, J. & Bader, G.D. Systematic analysis of somatic mutations in phosphorylation signaling predicts novel cancer drivers. Mol Syst Biol 9, 637 (2013).

25. Ryslik, G.A., Cheng, Y., Cheung, K.-H., Modis, Y. & Zhao, H. Utilizing protein structure to identify non-random somatic mutations. BMC Bioinformatics 14(2013).

26. Lahiry, P., Torkamani, A., Schork, N.J. & Hegele, R.A. Kinase mutations in human disease: interpreting genotype-phenotype relationships. Nat Rev Genet 11, 60–74 (2010).

27. Gross, A.M. et al. Multi-tiered genomic analysis of head and neck cancer ties TP53 mutation to 3p loss. Nat Genet 46, 939–43 (2014).

28. Jonsson, P.F. & Bates, P.A. Global topological features of cancer proteins in the human interactome. Bioinformatics 22, 2291–7 (2006).

29. AlQuraishi, M., Koytiger, G., Jenney, A., MacBeath, G. & Sorger, P.K. A multiscale statistical mechanical framework integrates biophysical and genomic data to assemble cancer networks. Nat Genet 46, 1363–71 (2014).

30. Nishi, H. et al. Cancer missense mutations alter binding properties of proteins and their interaction networks. PLoS One 8, e66273 (2013).

31. Cancer Genome Atlas Research, N. et al. The Cancer Genome Atlas Pan-Cancer analysis project. Nat Genet 45, 1113–20 (2013).

32. Lawrence, M.S. et al. Mutational heterogeneity in cancer and the search for new cancer-associated genes. Nature 499, 214–8 (2013).

33. Brooks, A.N., Kilgour, E. & Smith, P.D. Molecular pathways: fibroblast growth factor signaling: a new therapeutic opportunity in cancer. Clin Cancer Res 18, 1855–62 (2012).

34. Olson, M.F., Ashworth, A. & Hall, A. An essential role for Rho, Rac, and Cdc42 GTPases in cell cycle progression through G1. Science 269, 1270–1272 (1995).

35. Rooney, M.S., Shukla, S.A., Wu, C.J., Getz, G. & Hacohen, N. Molecular and genetic properties of tumors associated with local immune cytolytic activity. Cell 160, 48–61 (2015).

36. Brown, S.D. et al. Neo-antigens predicted by tumor genome meta-analysis correlate with increased patient survival. Genome Res 24, 743–50 (2014).

37. Markiewski, M.M. et al. Modulation of the antitumor immune response by complement. Nat Immunol 9, 1225–35 (2008).

38. Janelle, V. et al. Transient complement inhibition promotes a tumor-specific immune response through the implication of natural killer cells. Cancer Immunol Res 2, 200–6 (2014).

39. Rolland, T. et al. A proteome-scale map of the human interactome network. Cell 159, 1212–26 (2014).

40. Li, J. et al. TCPA: a resource for cancer functional proteomics data. Nat Methods 10, 1046–7 (2013).

41. Arkhipov, A. et al. Architecture and membrane interactions of the EGF receptor. Cell 152, 557–69 (2013).

42. Hao, Y. et al. Gain of interaction with IRS1 by p110alpha-helical domain mutants is crucial for their oncogenic functions. Cancer Cell 23, 583–93 (2013).

43. Follis, A.V. et al. The DNA-binding domain mediates both nuclear and cytosolic functions of p53. Nat Struct Mol Biol 21, 535–43 (2014).

44. Nehrt, N.L., Peterson, T.A., Park, D. & Kann, M.G. Domain landscapes of somatic mutations in cancer. BMC Genomics 13 Suppl 4, S9 (2012).

45. Porta-Pardo, E., Hrabe, T. & Godzik, A. Cancer3D: understanding cancer mutations through protein structures. Nucleic Acids Res (2014).

46. Supek, F., Minana, B., Valcarcel, J., Gabaldon, T. & Lehner, B. Synonymous mutations frequently act as driver mutations in human cancers. Cell 156, 1324–35 (2014).

47. Jaroszewski, L., Li, Z., Cai, X.H., Weber, C. & Godzik, A. FFAS server: novel features and applications. Nucleic Acids Res 39, W38–44 (2011).

48. Xu, D., Jaroszewski, L., Li, Z. & Godzik, A. AIDA: ab initio domain assembly server. NucleicAcids Res 42, W308–13 (2014).

49. Porta-Pardo, E. & Godzik, A. Analysis of individual protein regions provides novel insights on cancer pharmacogenomics. PLoS Comput Biol 11, e1004024 (2015).

50. Zhong, Q. et al. Edgetic perturbation models of human inherited disorders. Mol Syst Biol 5, 321 (2009).

51. Sahni, N. et al. Edgotype: a fundamental link between genotype and phenotype. Curr Opin Genet Dev 23, 649–57 (2013).

52. Ryan, C.J. et al. High-resolution network biology: connecting sequence with function. Nat Rev Genet 14, 865–79 (2013).

53. Wang, X. et al. Three-dimensional reconstruction of protein networks provides insight into human genetic disease. Nat Biotechnol 30, 159–64 (2012).

54. Garassino, M.C. et al. Different types of K-Ras mutations could affect drug sensitivity and tumour behaviour in non-small-cell lung cancer. Annals of Oncology 22(2011).

55. Ryan, C.J., Lord, C.J. & Ashworth, A. DAISY: picking synthetic lethals from cancer genomes. Cancer Cell 26, 306–8 (2014).

56. McLaren, W. et al. Deriving the consequences of genomic variants with the Ensembl API and SNP Effect Predictor. Bioinformatics 26, 2069–70 (2010).

57. Shoemaker, B.A. et al. Inferred Biomolecular Interaction Server--a web server to analyze and predict protein interacting partners and binding sites. Nucleic Acids Res 38, D518–24 (2010).

